# Maternal Odor Reduces the Neural Response to Fearful Faces in Human Infants

**DOI:** 10.1101/827626

**Authors:** Sarah Jessen

## Abstract

Maternal odor is known to play an important role in mother-infant-interaction in many altricial species such as rodents. However, we only know very little about its role in early human development. The present study therefore investigated the impact of maternal odor on infant brain responses to emotional expression. We recorded the electroencephalographic (EEG) signal of seven-month-old infants watching happy and fearful faces. Infants in two control groups exposed to no specific odor (control 1) or the odor of a different infant’s mother (control 2) showed the expected EEG fear response. Crucially, this response was markedly absent in the experimental group exposed to their mother’s odor. Thus, infants respond differently to fear signals in the presence of maternal odor. Our data therefore suggest that maternal odor can be a strong modulator of social perception in human infants.

## Introduction

As members of an altricial species, newborn humans completely rely on their social environment for survival. To foster and support the care they receive, newborns show a number of mechanisms to support social bonding, including a strong preference for faces (Johnson, Dziurawiec, Ellis, & Morton, 1991) and their mother’s voice (DeCasper & Fifer, 1980). However, face and voice are not the only sources of social information, and prior work suggests that olfaction and especially maternal odor can play an important role in early social development (Lubke & Pause, 2015).

One area in which the role of maternal odor has been amply investigated is breastfeeding. Human neonates respond to the smell of breast milk within days after birth (Doucet, Soussignan, Sagot, & Schaal, 2007; Marlier & Schaal, 2005; Porter, Makin, Davis, & Christensen, 1992), they prefer their mother’s unwashed over their mother’s washed breast (Varendi, Porter, & Winberg, 1994), and they quickly develop a preference for their own mother’s breast milk (Russell, 1976). Interestingly, maternal odor not only appears to facilitate nursing, but also seems to have a regulatory influence on other aspects of a neonate’s life. Maternal odor can have a soothing effect on crying infants (Sullivan & Toubas, 1998) and appears to reduce the pain response during medical procedures such as heel sticks (Nishitani et al., 2009; Zhang, Su, Li, & Chen, 2018).

Over the course of infancy, maternal odor can furthermore impact cognitive and perceptual processes. Importantly, the presence of maternal odor has been shown to impact face processing (Durand, Baudouin, Lewkowicz, Goubet, & Schaal, 2013; Durand, Schaal, Goubet, Lewkowicz, & Baudouin, 2020). Four-month-old infants tend to look longer at faces, and in particular the eye region of faces, in the presence of maternal odor (Durand et al., 2013). In a recent study, Leleu and colleagues (Leleu et al., 2019) furthermore investigated the influence of maternal odor on the neural response to faces in 4-month-old infants, and found an enhanced face-related neural response in the presence of maternal odor. In sum, maternal odor therefore appears to impact face processing in infancy both on a neural and a behavioral level. Interestingly, this effect appears to be specific to facial (or potentially social) information, as no comparable effect was found for non-social control stimuli (Durand et al., 2013). Furthermore, maternal odor also influences infants’ looking behavior to familiar compared to unfamiliar faces (Durand et al., 2020), suggesting that maternal odor no only modulates the response to faces per se, but also influences the processing of facial information.

However, facial identity is not the only information infants (and adults) can glean from faces; another prominent type of information that can be extracted from facial information is someone’s emotional state. The processing of emotional expressions has been amply investigated in human infants, and one prominent finding is that by about 7 months of age, infants discriminate between different emotional facial expression (for review, see Grossmann, 2010; Leppänen & Nelson, 2009, 2012). In particular, infants start to show an attentional bias towards fearful expressions (Vaish, Grossmann, & Woodward, 2008), which can be seen both on a neural (Leppänen, Moulson, Vogel-Farley, & Nelson, 2007; Peltola, Leppänen, Mäki, & Hietanen, 2009) and a behavioral level (Leppänen et al., 2007; Miguel, McCormick, Westerlund, & Nelson, 2019; Peltola, Hietanen, Forssman, & Leppänen, 2013). At the same time, recent work suggests that this fear bias can be strongly influenced by secondary factors, such as infant temperament (Martinos, Matheson, & de Haan, 2012) and breastfeeding experience (Krol, Rajhans, Missana, & Grossmann, 2014).

Importantly, these factors are linked to the interplay between the infant and their social environment, providing initial evidence for a modulation by social factors. However, at the same time, all the above-mentioned components (infant temperament, breastfeeding experience) are often interpreted as stable factors relating to interindividual differences rather than factors that flexibly change in a particular situation. Maternal odor in contrast is a situation-dependent signal that can either be present or absent in a given setting. It is therefore unclear whether a situation-dependent factor such as maternal odor can also impact infants’ response to fear signals.

To address this question, we designed an experiment to investigate the impact of maternal odor on the neural response to fear signals in human infants. In an electroencephalographic (EEG) set-up, infants were presented with happy and fearful facial expressions while they were exposed to either the familiar maternal odor, to an unfamiliar mother’s odor, or to no specific odor at all. To quantify infants’ response to fear signals, we investigated the amplitude of the *Nc*, an infant event-related potential (ERP) component observed between 400 and 800 ms after the onset of a stimulus at frontocentral electrodes. The Nc amplitude has been linked to the allocation of attention (Conte, Richards, Guy, Xie, & Roberts, 2020; Riggins & Scott, 2019; Webb, Long, & Nelson, 2005) and is typically enhanced in response to fearful faces in 7-month-old infants (Peltola et al., 2009).

Since prior studies suggest that long-term social factors such as extended breastfeeding experience can be associated with bias towards positive rather than negative facial expressions (Krol et al., 2014), we expect a reduction in the infant fear response by short-term social factors such as the mother’s presence, even if this presence is only signaled via maternal odor. In contrast, we predict that infants show the typical increased response to fearful faces in the absence of their mother’s odor.

## Methods

### Participants

Seventy-six 7-month-old infants were included in the final sample (age: 213 ± 8 days [mean ± standard deviation (SD)]; range: 200-225 days; 38 female, see Table 1 for description of the individual groups). An additional 15 infants had been tested but were not included in the final sample because they did not provide at least 10 artifact-free trials per condition (n=11); had potential neurological problems (n=1); were erroneously invited too young (n=1); the mean ERP response in the time-window and electrodes of interest was more than 4 standard deviations from the mean (n=1, see below); or because of technical problems during the recording (n=1).

**Table 1.** Overview of participants included in the final analysis. An additional 15 infants were tested but not included in the final analysis for various reasons (see text).

|  | N | female | age<br>(in days)* | still<br>breastfed | trials<br>(happy)* | trials<br>(fearful)* | Inf Neg<br>Temp** | EPDS** |
| --- | --- | --- | --- | --- | --- | --- | --- | --- |
| Maternal odor | 25 | 13 | $213 \pm 7$ | 14 | $38 \pm 18$ | $38 \pm 17$ | $3.03 \pm 0.68$ | $4.32 \pm 3.74$ |
| No odor | 26 | 9 | $214 \pm 8$ | 18 | $45 \pm 21$ | $45 \pm 23$ | $3.34 \pm 0.68$ | $5.08 \pm 4.77$ |
| Stranger odor | 25 | 16 | $215 \pm 7$ | 13 | $37 \pm 18$ | $37 \pm 17$ | $3.25 \pm 0.76$ | $4.54 \pm 3.96$ |

\* mean ± standard deviation; \* excluding one participant in the Stranger odor group, who did not fill in the questionnaire; Inf Neg Temp = Infant Negative Temperament, see text; EPDS = Edinburgh Postpartum Depression Screening, see text

The sample size was determined by statistical considerations and practical conventions in the field. First, for practical considerations and the known high attrition rates in infant EEG studies, we had planned a priori to keep collecting data until 25 useable data sets per each of the three experimental manipulation groups were obtained. Second, as outlined in Albers & Lakens (2018), a smallest effect size of interest was critical here, as too small true effects sizes for odor manipulations would not be of practical or translational relevance. In the present study, a total sample size of n=75 in three groups, was thus powered with 80% or more to detect medium and large effects (i.e., Cohen’s d of 0.8 or larger) at a conventional type I error level of 5 %.

Infants were recruited via the maternity ward at the local University hospital (Universitätsklinikum Schleswig-Holstein), were born full-term (38–42 weeks gestational age), had a birth weight of at least 2500 g, and had no known neurological deficits. The study was conducted according to the Declaration of Helsinki, approved by the ethics committee at the University of Lübeck, and parents provided written informed consent.

### Stimulus

As emotional face stimuli, we used colored photographs of happy and fearful facial expressions by 6 actresses from the FACES database (Ebner, Riediger, & Lindenberger, 2010 [actress-ID 54, 63, 85, 90, 115, 173]). Photographs were cropped so that only the face was visible in an oval shape, and have successfully been used in prior studies to investigate processing of emotional faces in infancy (Jessen & Grossmann, 2015, 2017).

### Odor manipulation

Prior to a scheduled experimental recording, all infants’ mothers were given a white cotton t-shirt and instructed to wear this t-shirt for three nights in a row. The mother was asked to store the t-shirt in a provided zip-lock bag during the day, and use her normal shampoo, soap, deodorant etc. as usual but refrain from using new products. Before the t-shirt was given to the mother, it had been washed with the same detergent for all t-shirts.

For practical reasons, the t-shirts used in the *Stranger odor* group had to be stored in a freezer (−20 °C) in the laboratory to allow swapping them between different mother-infant-dyads. This was done since freezing has been shown to conserve odor (Lenochova, Roberts, & Havlicek, 2009). To furthermore avoid any potential confound due to freezing, we asked *all* mothers, irrespective of later group assignment, to store the t-shirt in their freezer at home in a zip-lock bag for at least one night after wearing the t-shirt for three nights. In the *Maternal odor* group, in three cases, this was not possible as the t-shirt only arrived three days prior to the appointment, and in two cases the mother did not report whether the t-shirt had been stored in the freezer. For the remaining 20 infants in the *Maternal odor* group, the t-shirt had been stored in the freezer for at least one night. In the *No odor* group, t-shirts were unworn and hence freezing was irrelevant for odor emission (but mothers followed the same instructions to preserve blindness to condition assignment). In the *Stranger odor* group, all t-shirts except one had been stored in the freezer for at least one night.

### Randomization

Infants were randomly assigned to either the *Maternal odor* group or one of the control groups (*No odor* group or *Stranger odor* group; Figure 1). As only constraint to fully random assignment, we monitored as the study proceeded that groups did not differ in gender, age, or breastfeeding experience. Infants in the *Maternal odor* group were administered the t-shirt previously worn by their mother during the experiment. Infants in the *No odor* group were administered an unworn t-shirt. Infants in the *Stranger odor* group were administered a t-shirt previously worn by the mother of one of the other infants. The t-shirt of their own mother was stored in a freezer to be used as a stimulus for a different infant in the *Stranger odor* group. Except in one case, both, parents and the experimenter administering the t-shirt, were blind to the group assignment.

**Figure 1.**
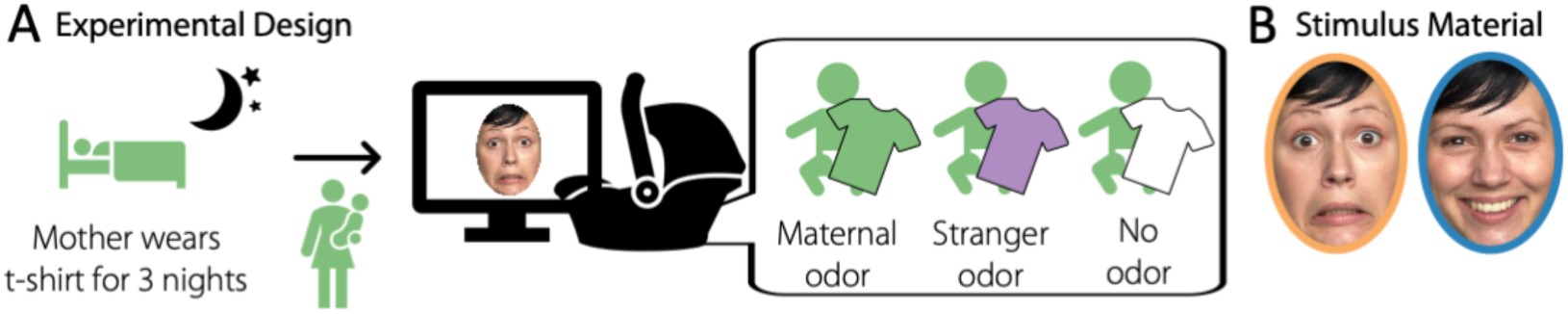
Experimental design. A) Mothers were asked to wear a provided t-shirt for 3 nights in a row prior to the experiment. The infant was randomly assigned to one of three groups; a Maternal odor group (exposed to the t-shirt worn by the infant’s mother), a Stranger odor group (exposed to a t-shirt worn by a different infant’s mother), or a No odor group (exposed to an unworn t-shirt). We recorded the EEG signal while the infants were seated in a car seat with the t-shirt positioned over their chest area and watched happy and fearful facial expressions. B) Example of fearful and happy faces used as stimulus material, the colored circles are for illustration purpose only and correspond to the color coding used in the following figures.

### Procedure and experimental design

Before the laboratory visit, families were sent the t-shirt (as described above) as well as a set of questionnaires, in particular the EPDS (Cox, Holden, & Sagovsky, 1987), the IBQ-R (Gartstein & Rothbart, 2003; Vonderlin, Ropeter, & Pauen, 2012), and a lab-internal questionnaire assessing demographic information as well as feeding and sleeping routines of the infant (One family, whose infant was assigned to the *Stranger odor* group, did not fill in the IBQ-R and the EPDS and is therefore not included in the control analyses with these two factors). After arriving in the laboratory, parents and infant were familiarized with the environment and parents were informed about the study and signed a consent form. The EEG recording was prepared while the infant was sitting on their parent’s lap. For recording, we used an elastic cap (BrainCap, Easycap GmbH) in which 27 AgAgCl-electrodes were mounted according to the international 10-20-system. An additional electrode was attached below the infant’s right eye to record the electrooculogram. The EEG signal was recorded with a sampling rate of 250 Hz using a BrainAmp amplifier and the BrainVision Recorder software (both Brain Products).

For the EEG recording, the infant was sitting in an age appropriate car seat (Maxi Cosi Pebble) positioned on the floor. The t-shirt was positioned over the chest area of the infant, folded along the vertical axis of the t-shirt and with the armpit region of the t-shirt directed towards the infant’s face. The t-shirt was fixated using the safety straps of the car seat as closely to the chin of the infant as possible and adjusted during the experiment if necessary.

In front of the infant (approximately 60 cm from the infant’s feet), a 24-inch monitor with a refresh rate of 60 Hz was positioned at a height of about 40 cm (bottom edge of the screen). The parent was seated approximately 1.5 m behind the infant and instructed not to interact with the infant during the experiment.

The experiment was programmed using the Presentation software (Version 18.1). Faces were presented for 800 ms, preceded by a fixation cross presented for 300 ms, and followed by an intertrial interval jittered between 800 and 1200 ms. The faces had a height of approximately 28 cm. If necessary, short video clips containing colorful moving shapes and ringtones were played during the experiment to redirect the infant’s attention to the screen. Each infant saw a maximum of 216 trials, arranged in miniblocks of 24 trials containing 12 happy and 12 fearful faces and played consecutively without interruption. Trials were presented in a pseudorandomized order, ensuring that no stimulus category (happy, fearful) was repeated more than once. The experiment continued until the infant had seen all trials or became too fussy to continue the experiment. During the experiment, the infant was video-recorded using a small camera mounted on top of the monitor to offline exclude trials in which the infant did not attend to the screen.

### Data Analysis

We analyzed the data using Matlab 2013b (The MathWorks, Inc., Natick, MA), the Matlab toolbox FieldTrip (Oostenveld, Fries, Maris, & Schoffelen, 2011), and for statistical analysis the package JASP (JASP Team, version 0.10.2).

### EEG Preprocessing

For purposes of artefact removal including an independent component analysis (ICA) routine, all data were first referenced to the average of all electrodes (average reference), filtered using a 100-Hz lowpass and a 1-Hz highpass filter, and segmented into 1-sec-epochs. To detect epochs obviously contaminated by artifacts, the standard deviation was computed in a sliding window of 200 msec. If the standard deviation exceeds 100 µV at any electrode, the entire epoch was discarded. Next, an independent component analysis (ICA) using the runica algorithm was computed on the remaining concatenated data.

Components were classified as artifactual based on visual inspection and rejected from the continuous, unfiltered data if classified as artefactual (4 ± 2 components per participants [mean ± SD], range 0–10 components).

After removal of ICA components, the data was re-segmented into epochs ranging from 200 ms before to 800 ms after the onset of the stimulus, re-referenced to the linked mastoids (mean of TP9 and TP10), and a 0.2 to 20 Hz bandpass filter was applied. A last step of automatic artifact detection was applied, rejecting all epochs in which the standard deviation exceeded 80 µV. Data was inspected visually for remaining artifacts, and all trials in which the infant did not attend to the screen (as assessed via the video recording during the experiment) were rejected (see Table 1 for number of remaining trials).

### ERP analysis

To analyze the Nc response, we computed the mean response in a time-window of 400–800 ms after stimulus onset across frontocentral electrodes (F3, Fz, F4, C3, Cz, C4; see Supplementary Material for an analysis of occipital electrodes, where no significant effect was found). The data in the 200 ms preceding the stimulus onset were used as baseline. One participant was rejected from further analysis because the difference in the mean response to happy and fearful faces in this time-window and electrode cluster was more than 4 standard deviations from the mean across all other participants. Mean responses were entered into a repeated measures ANOVA with the within-subject factor Emotion (happy, fear) and the between-subject factor Odor (maternal, stranger, no odor). Furthermore, we included the infant’s current breastfeeding status (whether s/he was still breastfed at the time of testing or not) as reported by the mother (Breastfed [yes,no]) as a covariate, as lactation may impact the mother’s body odor (McClintock et al., 2005). Student’s t-tests are computed as post-hoc tests and effect sizes are reported as partial eta squared (*η* _*p*_^*2*^) and Cohen’s *d*. In addition, we also performed the equivalent analysis using Bayesian statistics; BF _10_ values above 1 are interpreted as anecdotal evidence, above 3 as moderate evidence, and above 10 as strong evidence for the research hypothesis (Wagenmakers et al., 2018).

To further analyze the Emotion effect, we ran a cluster-based permutation test (Maris & Oostenveld, 2007). Importantly, such a test does not make any a priori assumptions regarding latency and topography of an effect, and therefore avoids potential biases due to selection of specific ERP components or time windows. We therefore chose to run this additional analysis to confirm the effects found in the more traditional ERP analysis. We ran the test with 1000 permutations contrasting responses to happy and fearful faces separately for each *Odor* group. A cluster had to comprise at least 2 adjacent electrodes, was computed across time and electrode position, and a type-1-error probability of less than 0.05 at the cluster-level was ensured.

### Negative Affect

Negative affect was computed as the mean of the IBQ-R scales Sadness, Fear, and Distress to Limitations (Aktar et al., 2018).

## Results

### Influence of maternal odor on the Nc response

As predicted, we observed an overall enhanced Nc amplitude in response to fearful faces (significant main effect of Emotion [*F*(1,72) = 11.60, *p* = .001, *η η* _*p*_^*2*^ = 0.14; BF _10_ = 2.578]). Most importantly, however, this emotion effect critically depended on the odor group an infant had been assigned to (significant interaction Emotion × Odor [*F*(2,72) = 5.57, *p* = .006, *η η* _*p*_^*2*^= 0.13; BF _10_ = 4.564; Figure 2]).

**Figure 2.**
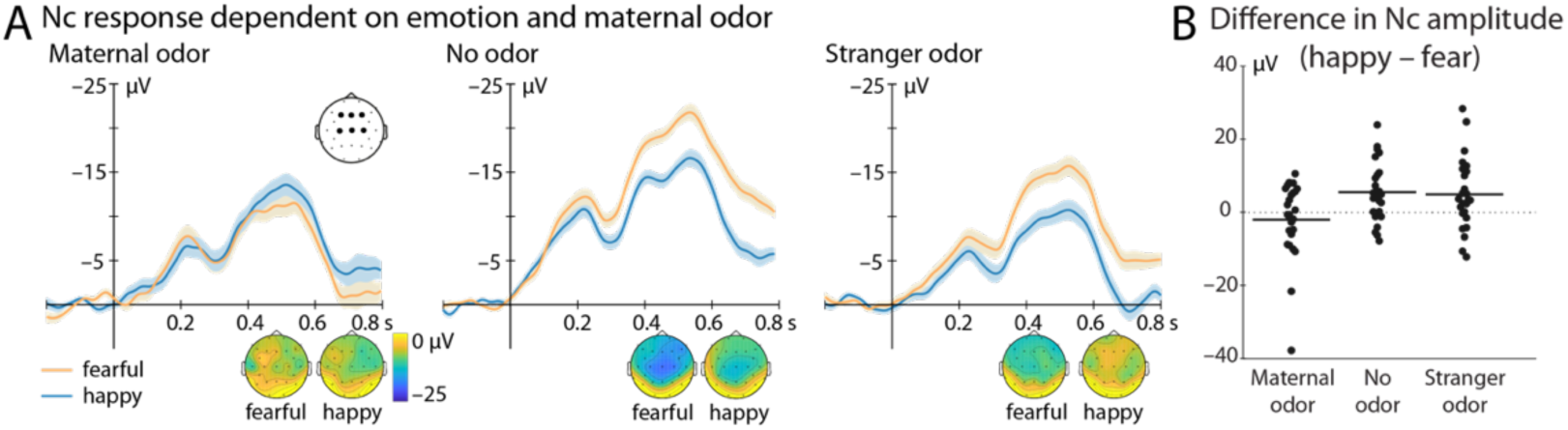
ERP response in the different odor groups. A) Shows the Nc response at frontocentral electrodes (F3, Fz, F4, C3, Cz, C4, marked by black dots) to fearful (orange) and happy (blue) facial expressions. While no difference in response was observed in the Maternal odor group, infants in the No odor and the Stranger odor group showed a significantly enhanced Nc response to fearful faces. Topographic representations averaged between 400 and 800 ms after face onset are shown at the bottom. B) Depicts the difference between Nc response to fearful and happy faces for each individual subject separately for the odor groups at the same electrodes and time window as in A. Mean difference is marked by horizontal black lines. Note that the interaction Odor × Emotion is significant even when excluding the two participants with the largest difference between happy and fear in the Maternal odor group.

Follow-up tests confirmed that the Nc effect to fearful faces was critically absent in the *Maternal odor* group [*t*(24) = –0.95, *p* = .35, *d* = –.19; BF _10_ = 0.32; fearful: –6.51 ± 2.99 µV, happy: –8.53 ± 3.17 µV]. In contrast, the typical enhancement of the Nc response to fearful (compared to happy) faces was present in the *Stranger odor* group [*t*(24) = 2.51, *p* = .019, *d* = .50; BF _10_ = 2.78; fearful: –10.57 ± 2.34 µV (mean ± SE), happy: –5.66 ± 1.78 µV] as well as in the *No odor* group [*t*(25) = 3.50, *p* = .002, *d* = .68; BF _10_ = 21.02; fearful: –16.94 ± 2.19 µV, happy: –11.43 ± 2.53 µV].

### Corroborating analysis using a cluster-based permutation approach

While the electrode and time window selection for this analysis had not been data derived but followed standards set by previous studies (Jessen & Grossmann, 2014, 2016, 2019), we aimed to corroborate this main result by a more data-driven search for potential effects using a cluster-based permutation test (Figure 3). In both, the *No odor* group and the *Stranger odor* group, nearly identical clusters indicating a significantly response enhancement to fearful (compared to happy) faces was found (No odor: *p* = .006, *T* _*sum*_ = 3063.8; Stranger odor: *p* = .021, *T* _*sum*_ = 1272.8). Importantly, both clusters exhibit the latency and topographic distribution typical for an Nc response. Most importantly, no such cluster of significant differences was found in the *Maternal odor* group when contrasting responses to happy and fearful faces.

**Figure 3.**
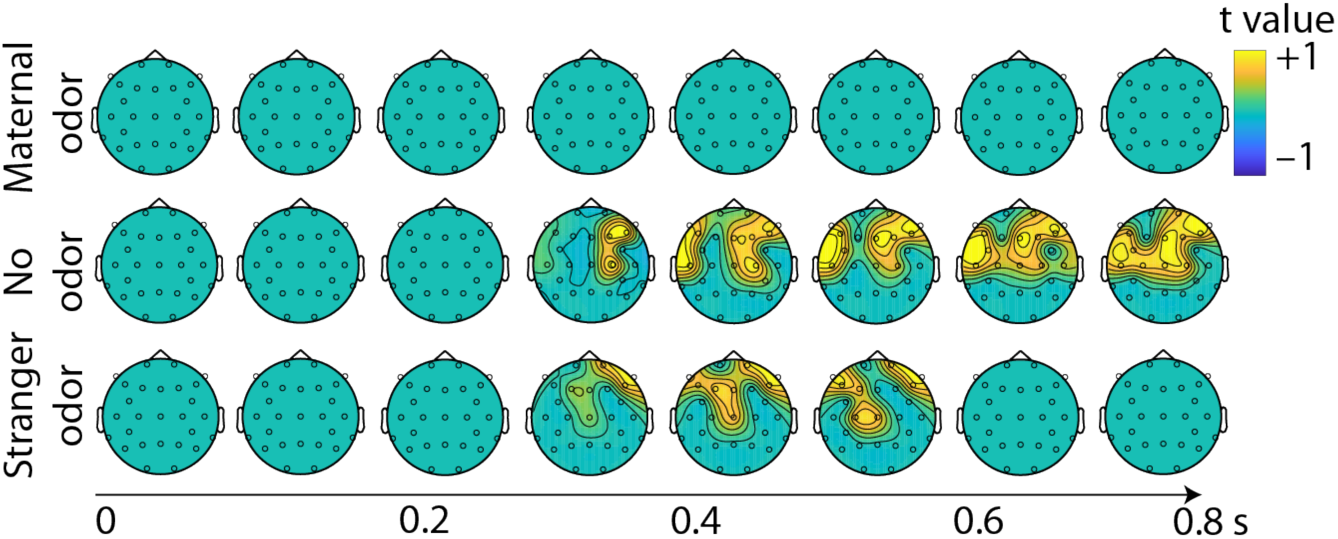
Cluster-based permutations test comparing responses to fearful and happy faces in the different odor groups. Depicted are topographic representations of t-values starting from the picture onset in steps of 100 ms.

Hence, while both control groups (*No group* and *Stranger odor* group) showed the age-typical enhanced Nc response to fearful faces, a heightened response to fearful faces was absent in the *Maternal odor* group. Our results suggest that maternal odor, as a signal of familiarity and maternal presence, reduces infant’s attention allocation to fear signals.

### No group differences with respect to potential confounds

Importantly, we did not find a difference between the three groups with respect to a number of potential confounds: There were no group differences in the number of included trials per infant in either Emotion condition [happy: *F*(2,73) = 1.49, *p* = .23, BF _10_ = 0.355 ; fearful: *F*(2,73) = 1.25, *p* = .29, BF _10_ = 0.296]; age [*F*(2,73) = 0.49, *p* = .61, BF _10_ = 0.165]; no differences in maternal depression scores as assessed via the EPDS [*F*(2,72) = 0.22, *p* = .80, BF _10_ = 0.136]; nor in infant negative temperament as assessed via the IBQ-R [*F*(2,72) = 1.23, *p* = .30, BF _10_ = 0.294].

### Effect of Breastfeeding

A last finding supported our general line of reasoning. Namely, we did observe an interaction between Nc response to the emotional expression of the presented face and whether the infant was still breastfed or not [Emotion × Breastfeeding, *F*(1,72) = 5.06, *p* = .028, *η η* _*p*_^*2*^ = 0.07; BF _10_ = 1.632; Figure 4]. Only the infants who were not breastfed any more at the time of testing showed an enhanced Nc response to fearful faces [*t*(30) = 3.55, *p* = .001, *d* = .64; BF _10_ = 26.54; fearful: –13.35 ± 2.18 µV, happy: –7.90 ± 2.00 µV], while this enhancement was absent in the infants who were still breastfed [*t*(44) = 0.65, *p* = .52, *d* = 0.1; BF _10_ = 0.20; fearful: –10.08 ± 2.08 µV, happy: –9.05 ± 2.10 µV].

**Figure 4.**
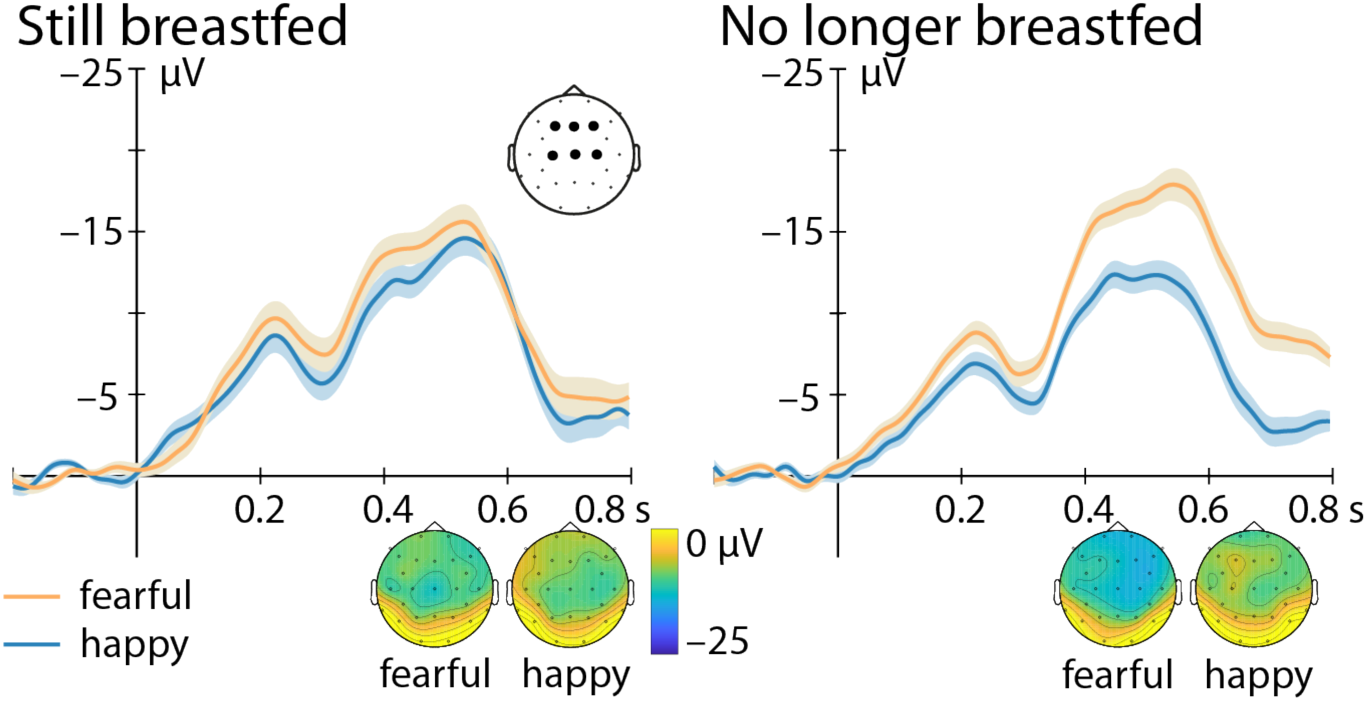
Nc response depending on breastfeeding status. Nc response is depicted at frontocentral electrodes (F3, Fz, F4, C3, Cz, C4, marked by black dots) to fearful (orange) and happy (blue) facial expressions for infants who are still breastfed (left) and not breastfed anymore (right). Of the infants not breastfed anymore, 9 had never been breastfed and the remaining 22 infants had been breastfed for some time (on average for 2.8 ± 2.3 months [mean ± standard deviation] after birth). Infants who are not breastfed any more show an enhanced Nc response to fearful faces, while this effect was absent in the group of infants who were still breastfed. Topographic representations averaged between 400 and 800 ms after face onset are shown at the bottom.

Importantly, this was independent of (i.e., additionally true but not interacting with) the *odor group* manipulation, as there was no meaningful Emotion × Breastfeeding × Odor interaction [*F*(2,70) = 2.20, *p* = .12, *η η* _*p*_^*2*^ = 0.06, BF _10_ = 1.081].

## Discussion

Our results demonstrate that maternal odor is a sufficiently strong signal to reduce the typically observed attentional response to fearful faces in 7-month-old infants. A highly consonant effect was found for breastfeeding, suggesting that not only momentary states but also longer-lasting effects related to maternal presence impact responses to fear signals in infants.

### Maternal odor as a momentary modulator of infants’ responses to signals of fear

We suggest that such a response pattern might be characteristic for a developing system that on the one hand needs to establish a close bonding to a caregiver, typically the mother, while on the other hand learning to respond to potential threat signals in the environment. This has been indirectly suggested by studies in older children (Gee et al., 2014) as well as rodent research (Landers & Sullivan, 2012). Extending these lines of research, our findings provide evidence for flexible processing of fear signals depending on maternal odor in early human development.

One potential interpretation of the observed pattern might be that a diminished response to threat signals in maternal presence (indicated via maternal odor) could facilitate bonding. Following this line of reasoning, a positive evaluation of information and less attention to potential negative signals may increase positive affect towards the caregiver even in the presence of negative signals. In addition, if maternal presence works as a “safety signal”, requiring the infant to allocate less attention to negative signals, this might also free cognitive capacities in the infant for other processes, akin to previously reported improved cognitive performance in rat pups in the presence of familiar odor (Wigal, Kucharski, & Spear, 1984).

Our results further underscore the importance of odor in early social development. Three recent studies have suggested a modulation of infant face processing in general by the presence of maternal odor (Durand et al., 2013, 2020; Leleu et al., 2019). Most importantly, Leleu et al (2019) found an enhanced neural response to faces in the presence of maternal odor. While their work thereby shows a modulation by maternal odor of face processing per se, the present result suggest that maternal odor can furthermore impact neural responses to specific aspects of face processing. Maternal odor might therefore be an important guiding factor in emotional learning in infancy.

Specifically, we found an impact on the attention-related Nc component (Conte et al., 2020; Webb et al., 2005) but no influence on early visual processing (see supplementary material) or on the number of trials the infants watched. Therefore, we found no evidence for a general impact of maternal odor on sensory processing or compliance with the experiment, but rather odor specifically impacted the evaluation of facial information, further underscoring its potential role in early social learning.

Importantly, the present manipulation did not differentiate between body odor and other odor components (such as deodorant used or specific food consumed by the mother), thereby reflecting the mélange of odors the infant experiences in maternal presence in everyday life. Hence, with the present approach, we cannot assess whether the observed effect can be attributed to the mother’s genuine body odor or rather to the overall familiar odor of the mother and the home environment. An extension of the present work separating these two potential sources – maternal body odor and overall familiar odor – may therefore provide interesting insights into the specificity of the current effect.

A further interesting factor in this context is parental proximity. As the infant grows more independent, detecting and responding to potential threat becomes of growing importance, especially if the mother is not present. At the same time, odor is a signal that is closely linked to parental proximity and/or familiar environment, hence the role of maternal odor during this period might be particularly interesting. Crucially, 7 months is an important turning point in early human development, characterized not only by qualitative changes in emotion development, but also by the onset of locomotion, an important step towards growing independence (Leppänen & Nelson, 2012). During this period, flexible responses to potential threats might be of particular importance, akin to what has been suggested in the rodent literature (Landers & Sullivan, 2012).

One important difference between the present study and most prior work on infant emotion perception is the positioning of the infant during the experiment; while the infants in the present study were seated in a car seat about 1.5 m apart from the parent, most other studies investigating infant emotion perception record data while the infant is sitting on their parent’s lap, hence in direct physical contact with the parent (e.g., Jessen & Grossmann, 2015; Leppänen et al., 2007; Xie, McCormick, Westerlund, Bowman, & Nelson, 2018). It might therefore be of interest in future studies to systematically manipulate parental proximity, its potential impact on infant responses to emotional signals and on the role of maternal odor.

### Breastfeeding as a long-term modulator of infants’ responses to signals of fear

While maternal odor as a situation-dependent or phasic signal influenced infants’ responses to fearful faces, so did the more tonic variable of an infant’s breastfeeding experience. Infants who were not breastfed any more at the time of the experiment did show the expected enhanced Nc response to fearful faces, while this was not the case for the infants who were still breastfed. These findings are in line with prior studies reporting an increased bias towards expressions of happiness with increasing breastfeeding experience (Krol, Monakhov, Lai, Ebstein, & Grossmann, 2015; Krol et al., 2014). How exactly breastfeeding experience interacts with emotion processing is not certain, but a possible explanation is an increased closeness between mother and infants; breastfed infants on average spend more time interacting with their mother (Smith & Forrester, 2017) and show a higher attachment security (Gibbs, Forste, & Lybbert, 2018). However, such reasoning would go against prior work suggesting that an enhanced fear response at seven months is indicative of better attachment quality (Peltola, Forssman, Puura, van Ijzendoorn, & Leppänen, 2015; Peltola, van IJzendoorn, & Yrttiaho, 2020). Hence, future studies systematically discerning breastfeeding experience from other variables related to mother-infant-interaction should assess the implications of this effect for socioemotional development.

In sum, our findings extend prior research suggesting an impact of breastfeeding experience on emotion processing in infancy. Factors related to maternal presence may therefore not only modulate responses to fearful faces directly, as suggested by the influence of maternal odor, but might also exert a longer-lasting impact.

### Future Directions and Limitations

While the present findings provide first evidence for an impact of maternal odor on emotion perception in infancy, future studies are clearly needed to further characterize the role its role in early social processing. Two important factors for future studies already mentioned are discerning maternal body odor from other types of familiar odor and the role of parental proximity (see above).

Another important aspect are potential changes across development. In rodents, it has been suggested that maternal presence, which can be signaled by maternal odor, may have a modulatory effect on offspring fear learning, in particular during the period in development when the offspring starts to spend increasing amounts of time away from their mother (for review, see Landers & Sullivan, 2012). One interesting approach for future studies is therefore the question whether a similar pattern can be observed in humans: is there a specific time-window during which infants show flexible responses to fear signals depending on the presence, and by extension the odor, of their mother?

Interestingly, a prime candidate for such a time window might be around seven months of age, when infants not only start to discriminate emotional expressions but also for the first time acquire the ability to locomote (see e.g. Campos et al., 2000; Leppänen & Nelson, 2012). At the same time, while most studies report an onset of the fear-bias between 5 and 7 months of age, several recent studies point to a potential earlier onset (e.g. Bayet et al., 2017; Heck, Hock, White, Jubran, & Bhatt, 2016; Safar & Moulson, 2020). Furthermore, prior studies showing an impact of maternal odor on face processing investigated infants at 4 months of age (Durand et al., 2013, 2020; Leleu et al., 2019), showing that maternal odor influences face processing per se already at an earlier age than investigated here. Hence, tracing the impact of maternal odor on emotional face processing longitudinally may be a promising approach to further assess the interplay between both factors.

Finally, the generalizability to other types of signals needs to be assessed in future work. We show that maternal odor influences the age-typical attentional response to fearful faces (as indicated via the Nc response), which constitute a particular instance of negative social information. The first question that arises is whether maternal odor also impacts infants’ responses to other negative but not necessarily social signals, such as pain or aversive sounds. Since recent studies suggest a link between maternal odor and the processing of faces in infancy (Durand et al., 2013; Leleu et al., 2019), one could also expect that this effect may be specific to social compared to non-social types of information.

At the same time, recent findings show that maternal odor can also impact the processing of facial identity (Durand et al., 2020), suggesting that maternal odor might impact different aspects of face processing beyond responses to facial emotional expressions. Future studies are needed to assess the robustness of the present findings in larger samples, and to test the generalizability to different types of social and non-social signals.

### Conclusions

The current study demonstrates that maternal odor influences the brain response to fearful facial expressions in infancy. While infants in two control groups of different specificity (a different mother’s odor or no specific odor at all) showed an expectably enhanced attentional response to fear signals (as indicated via the Nc amplitude), this response was absent in infants who could smell their mother. Our results establish that the mother’s presence, even if just signaled by the mother’s familiar odor, can result in a marked reduction of the neurobiological response to fear signals in infants. Furthermore, our data provide evidence for the potency of odor as a social signal in humans and in particular in early ontogeny.

## Acknowledgements

This work was supported by funding of the German Research Foundation (DFG, grant-number JE 781/1-1 & 2). We thank Leonie Emmerich and Aylin Ulubas for help with the data acquisition, Jonas Obleser for helpful comments on the manuscript, and all the families for participating.

## Bibliography

Aktar, E., Mandell, D. J., de Vente, W., Majdandžić, M., Oort, F. J., van Renswoude, D. R., … Bögels, S. M. (2018). Parental negative emotions are related to behavioral and pupillary correlates of infants’ attention to facial expressions of emotion. Infant Behavior and Development. https://doi.org/10.1016/j.infbeh.2018.07.004

Albers, C., & Lakens, D. (2018). When power analyses based on pilot data are biased: Inaccurate effect size estimators and follow-up bias. Journal of Experimental Social Psychology. https://doi.org/10.1016/j.jesp.2017.09.004

Bayet, L., Quinn, P. C., Laboissière, R., Caldara, R., Lee, K., & Pascalis, O. (2017). Fearful but not happy expressions boost face detection in human infants. Proceedings of the Royal Society B: Biological Sciences. https://doi.org/10.1098/rspb.2017.1054

Campos, J. J., Anderson, D. I., Barbu-Roth, M. A., Hubbard, E. M., Hertenstein, M. J., & Witherington, D. (2000). Travel broadens the mind. Infancy, 1(2), 149–219.

Conte, S., Richards, J. E., Guy, M. W., Xie, W., & Roberts, J. E. (2020). Face-sensitive brain responses in the first year of life. NeuroImage. https://doi.org/10.1016/j.neuroimage.2020.116602

Cox, J. L., Holden, J. M., & Sagovsky, R. (1987). Detection of postnatal depression. Development of the 10-item Edinburgh Postnatal Depression Scale. British Journal of Psychiatry, 150, 782–786. Retrieved from http://www.ncbi.nlm.nih.gov/pubmed/3651732

DeCasper, A. J., & Fifer, W. P. (1980). Of human bonding: newborns prefer their mothers’ voices. Science, 208(4448), 1174–1176. Retrieved from http://www.ncbi.nlm.nih.gov/pubmed/7375928

Doucet, S., Soussignan, R., Sagot, P., & Schaal, B. (2007). The “smellscape” of mother’s breast: Effects of odor masking and selective unmasking on neonatal arousal, oral, and visual responses. Developmental Psychobiology. https://doi.org/10.1002/dev.20210

Durand, K., Baudouin, J. Y., Lewkowicz, D. J., Goubet, N., & Schaal, B. (2013). Eye-Catching Odors: Olfaction Elicits Sustained Gazing to Faces and Eyes in 4-Month-Old Infants. PLoS ONE. https://doi.org/10.1371/journal.pone.0070677

Durand, K., Schaal, B., Goubet, N., Lewkowicz, D. J., & Baudouin, J. Y. (2020). Does any mother’s body odor stimulate interest in mother’s face in 4-month-old infants? Infancy. https://doi.org/10.1111/infa.12322

Ebner, N. C., Riediger, M., & Lindenberger, U. (2010). FACES--a database of facial expressions in young, middle-aged, and older women and men: development and validation. Behavior Research Methods, 42(1), 351–362. https://doi.org/10.3758/BRM.42.1.351

Gartstein, M. A., & Rothbart, M. K. (2003). Studying infant temperament via the Revised Infant Behavior Questionnaire. Infant Behavior and Development, 26(1), 64–86.

Gee, D. G., Gabard-Durnam, L., Telzer, E. H., Humphreys, K. L., Goff, B., Shapiro, M., … Tottenham, N. (2014). Maternal Buffering of Human Amygdala-Prefrontal Circuitry During Childhood but Not During Adolescence. Psychological Science. https://doi.org/10.1177/0956797614550878

Gibbs, B. G., Forste, R., & Lybbert, E. (2018). Breastfeeding, Parenting, and Infant Attachment Behaviors. Maternal and Child Health Journal. https://doi.org/10.1007/s10995-018-2427-z

Grossmann, T. (2010). The development of emotion perception in face and voice during infancy. Restorative Neurology and Neuroscience, 28, 236–291.

Heck, A., Hock, A., White, H., Jubran, R., & Bhatt, R. S. (2016). The development of attention to dynamic facial emotions. Journal of Experimental Child Psychology, 147, 100–110. https://doi.org/10.1016/j.jecp.2016.03.005

Jessen, S., & Grossmann, T. (2014). Unconscious discrimination of social cues from eye whites in infants. Proceedings of the National Academy of Sciences of the United States of America, 111(45), 16208–16213. https://doi.org/10.1073/pnas.1411333111

Jessen, S., & Grossmann, T. (2015). Neural signatures of conscious and unconscious emotional face processing in human infants. Cortex, 64, 260–270. https://doi.org/10.1016/j.cortex.2014.11.007

Jessen, S., & Grossmann, T. (2016). The developmental emergence of unconscious fear processing from eyes during infancy. Journal of Experimental Child Psychology, 142, 334–343. https://doi.org/10.1016/j.jecp.2015.09.009

Jessen, S., & Grossmann, T. (2017). Exploring the role of spatial frequency information during neural emotion processing in human infants. Frontiers in Human Neuroscience, 11. https://doi.org/10.3389/fnhum.2017.00486

Jessen, S., & Grossmann, T. (2019). Neural evidence for the subliminal processing of facial trustworthiness in infancy. Neuropsychologia, 126, 46–53. https://doi.org/10.1016/j.neuropsychologia.2017.04.025

Johnson, M. H., Dziurawiec, S., Ellis, H., & Morton, J. (1991). Newborns’ preferential tracking of face-like stimuli and its subsequent decline. Cognition, 40(1–2), 1–19. Retrieved from http://www.ncbi.nlm.nih.gov/pubmed/1786670

Krol, K. M., Monakhov, M., Lai, P. S., Ebstein, R. P., & Grossmann, T. (2015). Genetic variation in CD38 and breastfeeding experience interact to impact infants’ attention to social eye cues. Proceedings of the National Academy of Sciences of the United States of America, 112(39), E5434–42. https://doi.org/10.1073/pnas.1506352112

Krol, K. M., Rajhans, P., Missana, M., & Grossmann, T. (2014). Duration of exclusive breastfeeding is associated with differences in infants’ brain responses to emotional body expressions. Frontiers in Behavioral Neuroscience, 8, 459. https://doi.org/10.3389/fnbeh.2014.00459

Landers, M. S., & Sullivan, R. M. (2012). The development and neurobiology of infant attachment and fear. Developmental Neuroscience, 34(2–3), 101–114. https://doi.org/000336732

Leleu, A., Rekow, D., Poncet, F., Schaal, B., Durand, K., Rossion, B., & Baudouin, J.-Y. (2019). Maternal odor shapes rapid face categorization in the infant brain. Dev Sci. Retrieved from https://doi.org/10.1111/desc.12877

Lenochova, P., Roberts, S. C., & Havlicek, J. (2009). Methods of human body odor sampling: the effect of freezing. Chemical Senses, 34(2), 127–138. https://doi.org/10.1093/chemse/bjn067

Leppänen, J. M., Moulson, M. C., Vogel-Farley, V. K., & Nelson, C. A. (2007). An ERP study of emotional face processing in the adult and infant brain. Child Development, 78(1), 232–245. https://doi.org/10.1111/j.1467-8624.2007.00994.x

Leppänen, J. M., & Nelson, C. A. (2009). Tuning the developing brain to social signals of emotions. Nature Reviews: Neuroscience, 10, 37–47.

Leppänen, J. M., & Nelson, C. A. (2012). Early development of fear processing. Current Directions in Psychological Science, 21, 200–204.

Lubke, K. T., & Pause, B. M. (2015). Always follow your nose: the functional significance of social chemosignals in human reproduction and survival. Hormones and Behavior, 68, 134–144. https://doi.org/10.1016/j.yhbeh.2014.10.001

Maris, E., & Oostenveld, R. (2007). Nonparametric statistical testing of EEG- and MEG-data. Journal of Neuroscience Methods, 164(1), 177–190.

Marlier, L., & Schaal, B. (2005). Human newborns prefer human milk: Conspecific milk odor is attractive without postnatal exposure. Child Development. https://doi.org/10.1111/j.1467-8624.2005.00836.x

Martinos, M., Matheson, A., & de Haan, M. (2012). Links between infant temperament and neurophysiological measures of attention to happy and fearful faces. Journal of Child Psychology and Psychiatry and Allied Disciplines, 53(11), 1118–1127. https://doi.org/10.1111/j.1469-7610.2012.02599.x

McClintock, M. K., Bullivant, S., Jacob, S., Spencer, N., Zelano, B., & Ober, C. (2005). Human body scents: Conscious perceptions and biological effects. In Chemical Senses. https://doi.org/10.1093/chemse/bjh151

Miguel, H. O., McCormick, S. A., Westerlund, A., & Nelson, C. A. (2019). Rapid face processing for positive and negative emotions in 5-, 7-, and 12-month-old infants: An exploratory study. British Journal of Developmental Psychology. https://doi.org/10.1111/bjdp.12288

Nishitani, S., Miyamura, T., Tagawa, M., Sumi, M., Takase, R., doi, H., … Shinohara, K. (2009). The calming effect of a maternal breast milk odor on the human newborn infant. Neuroscience Research. https://doi.org/10.1016/j.neures.2008.10.007

Oostenveld, R., Fries, P., Maris, E., & Schoffelen, J.-M. (2011). FieldTrip: Open source software for advanced analysis of MEG, EEG, and invasive electrophysiological data. Computational Intelligence and Neuroscience, 2011, 156869.

Peltola, M. J., Forssman, L., Puura, K., van Ijzendoorn, M. H., & Leppänen, J. M. (2015). Attention to Faces Expressing Negative Emotion at 7 Months Predicts Attachment Security at 14 Months. Child Development. https://doi.org/10.1111/cdev.12380

Peltola, M. J., Hietanen, J. K., Forssman, L., & Leppänen, J. M. (2013). The emergence and stability of the attentional bias to fearful faces in infancy. Infancy, 18(6), 905–926.

Peltola, M. J., Leppänen, J. M., Mäki, S., & Hietanen, J. K. (2009). Emergence of enhanced attention to fearful faces between 5 and 7 months of age. Social Cognitive and Affective Neuroscience, 4, 134–142.

Peltola, M. J., van IJzendoorn, M. H., & Yrttiaho, S. (2020). Attachment security and cortical responses to fearful faces in infants. Attachment and Human Development. https://doi.org/10.1080/14616734.2018.1530684

Porter, R. H., Makin, J. W., Davis, L. B., & Christensen, K. M. (1992). Breast-fed infants respond to olfactory cues from their own mother and unfamiliar lactating females. Infant Behavior & Development, 15, 85–93.

Riggins, T., & Scott, L. S. (2019). P300 development from infancy to adolescence. Psychophysiology. https://doi.org/10.1111/psyp.13346

Russell, M. J. (1976). Human olfactory communication. Nature, 260(5551), 520–522. Retrieved from http://www.ncbi.nlm.nih.gov/pubmed/1264204

Safar, K., & Moulson, M. C. (2020). Three-month-old infants show enhanced behavioral and neural sensitivity to fearful faces. Developmental Cognitive Neuroscience. https://doi.org/10.1016/j.dcn.2020.100759

Smith, J. P., & Forrester, R. (2017). Maternal Time Use and Nurturing: Analysis of the Association between Breastfeeding Practice and Time Spent Interacting with Baby. Breastfeeding Medicine. https://doi.org/10.1089/bfm.2016.0118

Sullivan, R. M., & Toubas, P. (1998). Clinical usefulness of maternal odor in newborns: soothing and feeding preparatory responses. Biology of the Neonate, 74(6), 402–408. Retrieved from http://www.ncbi.nlm.nih.gov/pubmed/9784631

Vaish, A., Grossmann, T., & Woodward, A. (2008). Not all emotions are created equal: the negativity bias in social-emotional development. Psychological Bulletin, 134(3), 383–403. https://doi.org/10.1037/0033-2909.134.3.383

Varendi, H., Porter, R. H., & Winberg, J. (1994). Does the newborn baby find the nipple by smell? Lancet, 344(8928), 989–990. Retrieved from http://www.ncbi.nlm.nih.gov/pubmed/7934434

Vonderlin, E., Ropeter, A., & Pauen, S. (2012). Erfassung des fruehkindlichen Temperaments mit dem Infant Behavior Questionnaire Revised. Psychometrische Merkmale einer deutschen Version. Zeitschrift Fuer Kinder-Und Jugendpsychiatrie Und Psychotherapie, 40(5), 307–314.

Wagenmakers, E. J., Marsman, M., Jamil, T., Ly, A., Verhagen, J., Love, J., … Morey, R. D. (2018). Bayesian inference for psychology. Part I: Theoretical advantages and practical ramifications. Psychonomic Bulletin and Review. https://doi.org/10.3758/s13423-017-1343-3

Webb, S. J., Long, J. D., & Nelson, C. A. (2005). A longitudinal investigation of visual event-related potentials in the first year of life. Dev Sci, 8(6), 605–616. https://doi.org/10.1111/j.1467-7687.2005.00452.x

Wigal, T., Kucharski, D., & Spear, N. E. (1984). Familiar contextual odors promote discrimination learning in preweanling but not in older rats. Developmental Psychobiology. https://doi.org/10.1002/dev.420170512

Xie, W., McCormick, S. A., Westerlund, A., Bowman, L. C., & Nelson, C. A. (2018). Neural correlates of facial emotion processing in infancy. Developmental Science. https://doi.org/10.1111/desc.12758

Zhang, S., Su, F., Li, J., & Chen, W. (2018). The Analgesic Effects of Maternal Milk Odor on Newborns: A Meta-Analysis. Breastfeeding Medicine. https://doi.org/10.1089/bfm.2017.0226

