## Supplementary Material for "Maternal Odor Reduces the Neural Response to Fearful Faces in Human Infants"

*ERP analysis at occipital and parietal electrodes*

Although we did not have any a priori hypotheses regarding early visual processing, we also analyzed ERP responses at occipital electrodes (O1, O2; Figure S1), computing the same analysis as described for the Nc in the main text for the four time-windows 100 – 300 ms, 100 – 200 ms, 200 – 300 ms (covering the N290) and 300 – 500 ms (covering the P400) after stimulus onset based on visual inspection. We did not find any significant effect (see Table S1 for an overview of all results). In addition, we analyzed the N290 (200 – 300 ms) and the P400 (300 – 500 ms) in an extended ROI including both occipital (O1, O2) and parietal electrodes (P7, P8). Again, we did not find any significant effect (see Table S2). Note, however, that even if interpreting the interaction Emotion × Odor, this effect would not support the interpretation that Nc differences in emotion responses between the different Odor groups are driven by differences in early visual processing, since only the Stranger but not the No odor group show an early occipital difference, yet both show a robust difference at the Nc.

*Table S1. Overview of statistical comparisons at occipital electrodes (O1, O2).*

| | | |
|---|---|---|
| 100 – 300 ms | Emotion | $F(1,72) = 1.013, p = .318$ |
| | Emotion × Odor | $F(2,72) = 2.436, p = .095$ |
| | Emotion × Breastfeeding | $F(1,72) = 0.005, p = .942$ |
| 100 – 200 ms | Emotion | $F(1,72) = 3.072, p = .084$ |
| | Emotion × Odor | $F(2,72) = 2.587, p = .082$ |
| | Emotion × Breastfeeding | $F(1,72) = 0.030, p = .863$ |
| 200 – 300 ms | Emotion | $F(1,72) = 0.161, p = .689$ |
| | Emotion × Odor | $F(2,72) = 2.174, p = .121$ |
| | Emotion × Breastfeeding | $F(1,72) = 0.066, p = .798$ |
| 300 – 500 ms | Emotion | $F(1,72) = 1.165, p = .284$ |
| | Emotion × Odor | $F(2,72) = 1.218, p = .302$ |
| | Emotion × Breastfeeding | $F(1,72) = 1.301, p = .258$ |

*Table S2. Overview of statistical comparisons at occipital electrodes (O1, O2, P7, P8).*

| | | |
|---|---|---|
| 200 – 300 ms | Emotion | $F(1,72) = 0.510, p = .477$ |
| | Emotion × Odor | $F(2,72) = 2.259, p = .112$ |
| | Emotion × Breastfeeding | $F(1,72) = 0.072, p = .790$ |
| 300 – 500 ms | Emotion | $F(1,72) = 1.181, p = .281$ |
| | Emotion × Odor | $F(2,72) = 1.786, p = .175$ |
| | Emotion × Breastfeeding | $F(1,72) = 0.721, p = .399$ |

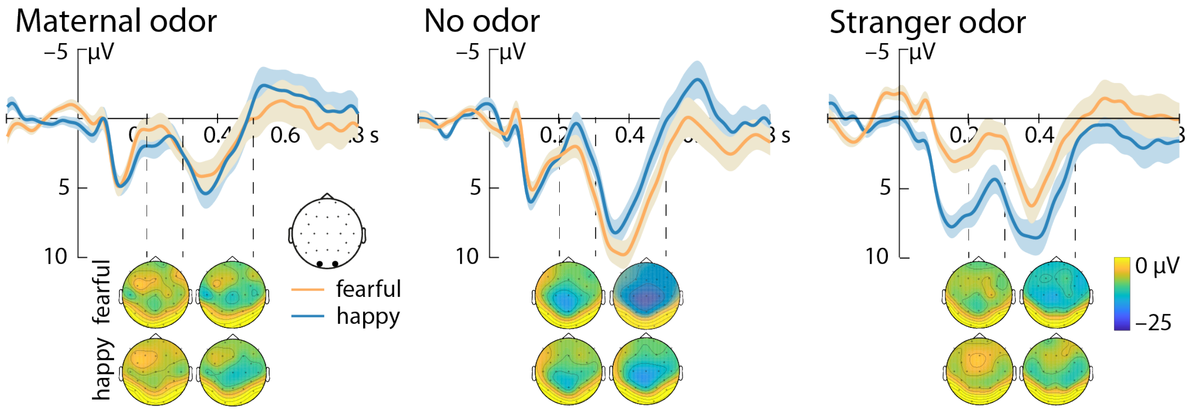


*Figure S1. ERP responses in the different odor groups at occipital electrodes (O1, O2). Responses to fearful faces are shown in orange, while responses to happy faces are shown in blue. Black dots on the topographic map indicate electrode position. Topographic representations are shown for the time-windows indicated by the dashed lines (200 – 300 ms and 300 – 500 ms), which correspond to the N290 and the P400, respectively.*
